# Combined optical coherence tomography and electroretinography (OCT+ERG) system for imaging neurovascular coupling in the human retina

**DOI:** 10.1101/2025.03.27.643714

**Authors:** Khushmeet Dhaliwal, Alexander Wong, Tom Wright, Kostadinka Bizheva

## Abstract

**Significance:** During their early stages of development, neurological and neurodegenerative diseases cause changes to the biological tissue’s morphology, physiology and metabolism at cellular level, and acute, transient changes in the local blood flow. Development of novel optical methods for quantitative imaging of such changes non-invasively and simultaneously would allow for probing of neurovascular coupling in neural tissues and therefore can have a profound effect on furthering our understanding of neurodegeneration.

**Aim:** To develop an optical imaging platform based on optical coherence tomography (OCT) for imaging and characterization of neurovascular coupling in the human retina with high spatial and temporal resolution.

**Approach:** A fast, ultrahigh resolution OCT system was developed and combined with a clinical electroretinography (ERG) system for in-vivo, simultaneous structural, functional and vascular imaging of the human retina in response to visual stimulation. Novel image processing algorithms were developed to quantify visually-evoked physiological and blood flow changes from the OCT images and explore neurovascular coupling in the healthy human retina.

**Results:** Visual stimulation of the human retina with singe flashes (white light, 4ms duration) caused transient changes in the optical reflectivity and thickness (optical pathlength difference) of major retinal layers, as well as the blood flow in local retinal blood vessels. The time courses of the neuronal and blood flow changes were correlated, and their magnitude was dependent on the intensity of the visual stimulus.

**Conclusions:** We have developed an optical imaging modality for non-invasive probing of neurovascular coupling in the living human retina and demonstrated its utility and clinical potential in a pilot study on healthy subjects. This imaging platform could serve as a useful clinical research tool for investigation of potentially blinding retinal diseases, as well as neurodegenerative brain diseases that are expressed in the retina such as Alzheimer’s and Parkinsons.

## 1 Introduction

The retina, often called ‘the window to the brain’, is an extension of the central nervous system (CNS) designed for detection of visual information and its transmission to the visual cortex of the brain.^1^ Retinal neurons of highly specialized function such as photoreceptors which detect light and filter it by spectral content, bipolar and horizontal cells which filter the photoreceptors responses spatially and temporally, and the ganglion cells which amplify the visual information and transmit it to the brain, are organized is well defined layers.^2–5^ Furthermore, the retina is highly vascularized with oxygen and nutrition delivered to the retinal neurons in the anterior retina by the inner retina vasculature, and to the photoreceptors by the choroidal vasculature.^6, 7^ Neurovascular coupling in the retina is referred to as a transient increase in the retinal blood flow and vasodilation / vasoconstriction of the retinal blood vessels in response to the increased metabolic demand of retinal neurons. This phenomenon was first observed in the brain^8^ and later, in the retina^9, 10^ using non-invasive optical imaging methods such as Laser Doppler Velocimetry.

Potentially blinding retinal diseases such Age-related Macular Degeneration (AMD),^11, 12^ Diabetic Retinopathy,^13–15^ Retinitis Pigmetosa^16–18^ and Glaucoma^19–21^ affect a large portion of the world’s aging population and amount to *>* $1.5B in direct costs to the Canadian Health Care system annually.^22^ These diseases cause progressive loss of normal function and eventually cell death of retinal neurons that in the later stages of disease the development are registered as morphological changes. In addition to causing metabolic and physiological changes in the retinal neurons, studies have reported blood flow changes with the progression of these retinal diseases.^23, 24^ However, the dynamic relationship between neuronal and blood flow changes (neurovascular coupling) is still not fully understood in the healthy and diseased human and animal retina partly due to the fact that functional and vascular studies have been conducted separately.

Optical Coherence Tomography (OCT) is an imaging modality that can generate non-invasively volumetric images of the retina with micrometer scale resolution, sufficient for visualization of the fine layered structure of the human retina.^25–29^ When combined with Adaptive Optics, OCT is also able to visualize individual retinal neurons.^30–32^ Doppler OCT can be used to measure retinal blood flow^33–35^ while optical microangiography (OMAG) can used to map the retinal vasculature in 3D over a wide field-of view.^36–38^ All of these OCT-based technologies have found valuable applications in ophthalmology for clinical research of various ophthalmic diseases.^18, 37, 39, 40^ Both Doppler OCT^41–43^ and OMAG^44, 45^ technologies have been used in the recent past to evaluate blood flow changes in the human retina evoked by visual stimulation. However, these studies utilized very long stimuli (> 5 seconds), had poor temporal resolution and did not record the retinal blood flow (RBF) changes continuously.^41, 43, 46, 47^ Therefore, they were not able to provide information of the transient behavior of the RBF with high temporal resolution.

Optoretinography (ORG) is the optical equivalent of electroretinography (ERG), which is a clinically established method for electrical recordings of the responses of retinal neurons to visual stimulation. The use of OCT for non-invasive, optical recording of visually evoked neuronal changes was first proposed by Bizheva and the proof-of-principle was demonstrated in in-vitro in animal tissue where OCT and ERG data were acquired synchronously.^48^ With advances in camera and laser technologies, over the past 20 years different OCT system designs were developed for ORG recordings of visually evoked neuronal activity in the human and animal retina: point scanning Spectral Domain OCT,^49^ AO-OCT,^30, 31^ Full-Field SS-OCT^50–53^ and Line-Field - SD-OCT.^54, 55^ While majority of the ORG data was acquired mostly from healthy retinas, most recent studies have explored changes in the ORG signal related to retinal diseases.^18^

Here we present the design of a novel, combined OCT+ERG system, novel imaging protocols and image processing algorithms that were developed for simultaneous probing of visually evoked changes in the human retina neuronal function and RBF with high spatial and temporalresolution. We demonstrated the utility of the OCT+ERG imaging modality as a new research tool for non-invasive probing of neurovascular coupling in the human retina.

## 2 Methods

The following paragraphs describe the optical design of the OCT system, the imaging protocols for the data collection and the image processing algorithms for analysis of the neuronal and vascular responses of the living retina to visual stimulation.

### 2.1 OCT+ERG system

A combined OCT+ERG system was developed for non-invasive probing of neurovascular coupling in the living human retina. The optical design of the compact, fiberoptic, spectral-domain OCT system is shown in Figure 1, while a photograph of the OCT imaging probe integrated with a custom visual stimulator is presented in Figure 1B. The output of a femtosecond laser (Femtolasers GmbH, Austria) was connected to a spool of a 100 m long fiber to stretch the femtosecond pulses and mimic CW emission over a broad spectral range (670 nm to 920 nm). The output from the spool was connected to a fiberoptic coupler (60*/*40 split ratio, Gould Fiberoptics, USA) that served as the core of the OCT interferometer.

**Fig 1.**
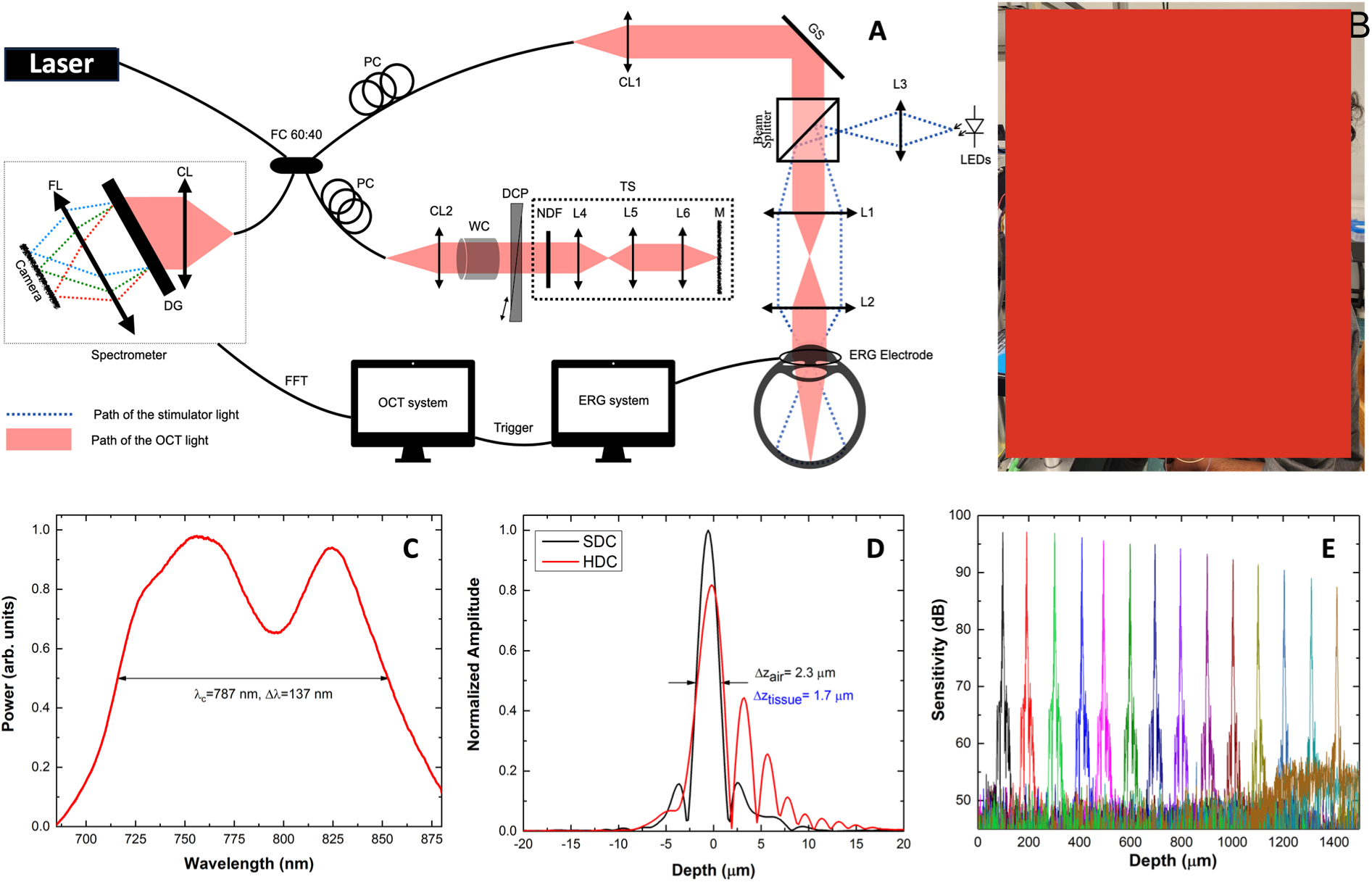
(A) Schematic of the combined OCT+ERG system. (B) Photograph of the OCT+ERG imaging probe. (C) Laser spectrum measured from the reference arm of the OCT system. (D) Axial OCT PSF after hardware (red color) and software (black color) dispersion compensation. (E) Sensitivity roll-off with imaging depth.

In the imaging arm of the OCT system, a parallel optical beam of 2.8 mm diameter was generated with a fiber collimator (CL1 - achromat doublet, f = 10 mm, Thorlabs, USA). A pair of galvanometric scanners (Cambridge Technologies, USA) were used to scan the imaging beam over the retinal surface. A telescope comprised of 2 achromat doublet lenses (L1 (f = 75 mm) and L2 (f = 60 mm), Thorlabs, USA) was used to generate a parallel optical beam of 2.24 mm diameter and optical power ∼ 1.1 mW that was projected onto the human cornea. The mechanical design of the telescope allowed for easy adjustment of the distance between lenses L1 and L2 to compensate for myopia and hyperopia in the imaged subjects and fine-tune the focusing of the imaging beam within the retina. A custom visual stimulator comprised of a wheel with 4 LEDs (white, blue, green and red) was integrated with the OCT imaging probe through lens L3 (f = 50 mm, Thorlabs, USA) and a pellicle beam splitter (T/R ratio of 92/8, Thorlabs, USA). This design allows for the light from one of the LEDs to be focused at the pupil plane of the human eye in order to generate wide field illumination of the retina with almost uniform intensity (Maxwellian view).

The custom visual stimulator was interfaced with a commercial electroretinography (ERG) system (Espion 2, Diagnosys LLC, USA), which served two purposes: a) precise control of the timing, duration, intensity and frequency of visual stimulus; and b) simultaneous acquisition of ERG and OCT recordings. The ERG recordings are used as a “gold standard” to validate the ORG traces extracted from the OCT images.

The reference arm of the OCT interferometer was comprised of a fiber collimator (achromat doublet, f = 10 mm, Thorlabs, USA), a custom-built 1 cm long water cell (WC) for crude compensation of water dispersion within the human eye, a custom dispersion compensation (DCP) unit comprised of BK7 prisms mounted on miniature translation stages (Edmund Optics, USA), a telescope (L4 and L5, achromat doublets (f4 = f5 = 60 mm), Thorlabs, USA), a focusing lens (L6, achromat doublet (f = 10 mm), Thorlabs, USA) and a mirror mounted on a miniature translation stage (Edmund Optics, USA). The telescope and the focusing mirror were mounted on a manual translation stage (Edmund Optics). Polarization controllers (PC) were used in both arms of the interferometer.

The detection end of the OCT system was comprised of a commercially available spectrometer (Cobra-S 800, Wasatch Photonics, Durham, USA), interfaced with a line-scan CMOS camera (OCTOPLUS, Teledyne, USA) with a readout rate of 250 kHz.

Figure 1C shows the spectrum of the optical beam measured at the detection end of the OCT system, which accounts for spectral changes induced by the optical transmission function of the OCT system. Figure 1D shows the axial PSF after hardware dispersion compensation only (red line) and after additional software dispersion compensation (black line), based on previous publications.^56, 57^ The OCT system’s maximum sensitivity was 98 dB with a roll-off of ∼ 10 dB over a scanning range of 1.5 mm, measured with 1.1 mW imaging power incident on the corneal surface which is well below the maximum permissible exposure (MPE) as specified by the ANSI standard (Figure 1E).

### 2.2 In-vivo imaging of the human retina

A pilot study to test the clinical viability of the OCT+ERG system was conducted on 2 healthy subjects (2 males, age range 25 yo ± 5 years). The study received full ethics clearance from the University of Waterloo’s Office of Research Ethics and a written consent was obtained from all participants. The study participants underwent standard clinical examination prior to the OCT imaging sessions and only subjects with no ocular diseases or a history of epilepsy and nearly perfect vision (minor correction up to -1 Diopters) were included in the study. A DTL type ERG electrode was inserted under the lower eyelid of the imaged eye, while reference and ground ERG electrodes are applied to the participant’s temple and behind the ear respectively (Figure 1B). Both pupils were dilated by application of 1 drop of Tropicamide in each eye followed by ∼ 30 minutes dark adaptation. A faint red LED was used as fixation target for the non-imaged eye.

### 2.3 Visual stimulation and image acquisition protocols

In this study, the retina was stimulated with white-light single flashes of 4 ms duration and different intensities ranging from 0*cd.s/m*^2^ (dark recordings) to 43.4*cd.s/m*^2^. OCT and ERG data were acquired over a period of ∼ 6 seconds (1 second pre-stimulus and 5 seconds post-stimulus), as shown in Figure 2A. Table 1 shows the percentage of photoreceptor bleaching for each stimulus intensity used in this study, calculated based on the method described by Rushton and Henry.^59, 60^

**Fig 2.**
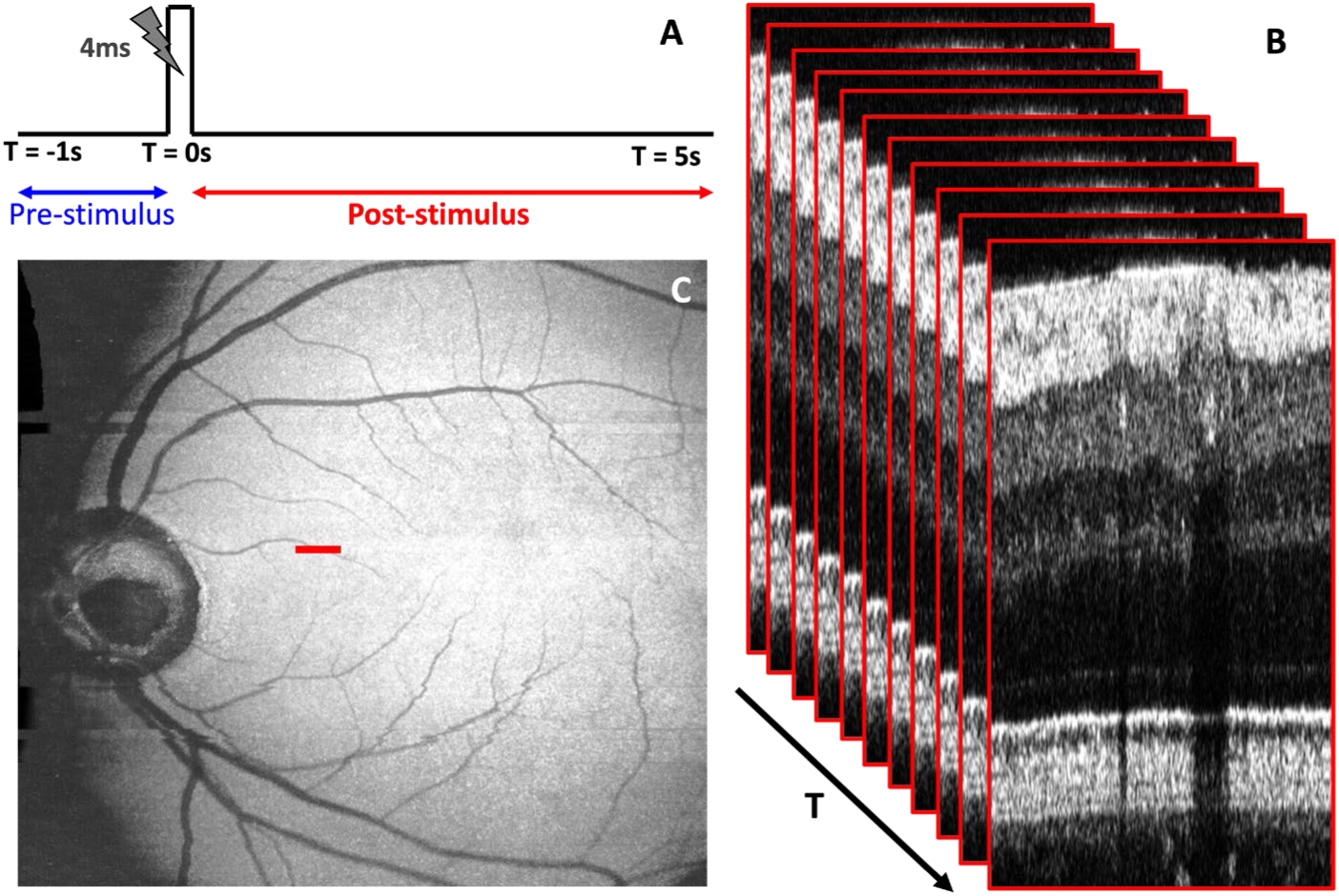
Image acquisition protocol for the optoretinogrpahy and functional blood flow recordings. (A) Timeline of the visual stimulus. (B) A sequence of 1,200 repeated B-scans acquired over duration of ∼ 6 seconds from a location in the retina approximately midway between the fovea and optic nerve head (C, red line).

**Table 1.** Bleaching percentage for different stimulus intensities.

| Stimulus Intensity<br>($cd.s/m^2$) | Bleaching<br>(%) |
| --- | --- |
| 1.4 | 1.4 |
| 14.5 | 13.8 |
| 43.4 | 36 |

Large field-of-view, volumetric morphological OCT images of the retina (1,000 A-scans × 1,000 B-scans) were acquired prior to acquisition of the functional OCT recordings, to map the vasculature on the retinal surface. Next, a series of 1,200 OCT B-scans, each composed of 1,000 A-scans (Figure 2B), were acquired from the same place in the retina located approximately 10*^◦^* from the fovea (red line in Figure 2C).

### 2.4 Image processing and data analysis

Custom MATLAB-based algorithms were developed for processing and analysis of the OCT data. The raw OCT data was first processed to generate dispersion-compensated complex-valued B-scans (representative intensity image shown in Figure 3A). Figure 3B shows a flow chart for the image processing steps designed to measure ORG responses (optical path length difference PD) and intensity changes) from different retinal layers. To account for involuntary eye motion caused by breathing, heartbeat and eye muscle twitching, a sub-pixel phase-restoring image registration method was utilized to correct for motion related misalignment between consecutive B-scans.^61–63^ Next, a region of interest (ROI) located outside the shadows of the retinal blood vessels was defined for each B-scan of the time series, the retinal layers were segmented automatically based on their intensity profiles using boundary detection^64^ and the B-scans were flattened using the photoreceptors IS-2 layer as a guideline. The segmented data was used to extract OPD and intensity changes for each retinal layer in response to visual stimulation. To generate the intensity traces, the values for all pixels within each segmented layer were averaged axially and standardizedintensity change was measured as described previously by Cooper *et. al.*^65, 66^ Optical path length changes within the segmented layers were calculated based on the following equation:

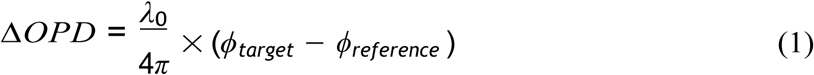

where, Δ*OPD* represents the optical path length difference changes between the reference and the target layers, *λ*_0_ is the central wavelength of the imaging light source, and (*ϕ_target_*− *ϕ_reference_*) represents the phase difference between the target retinal layer and the reference layer.

**Fig 3.**
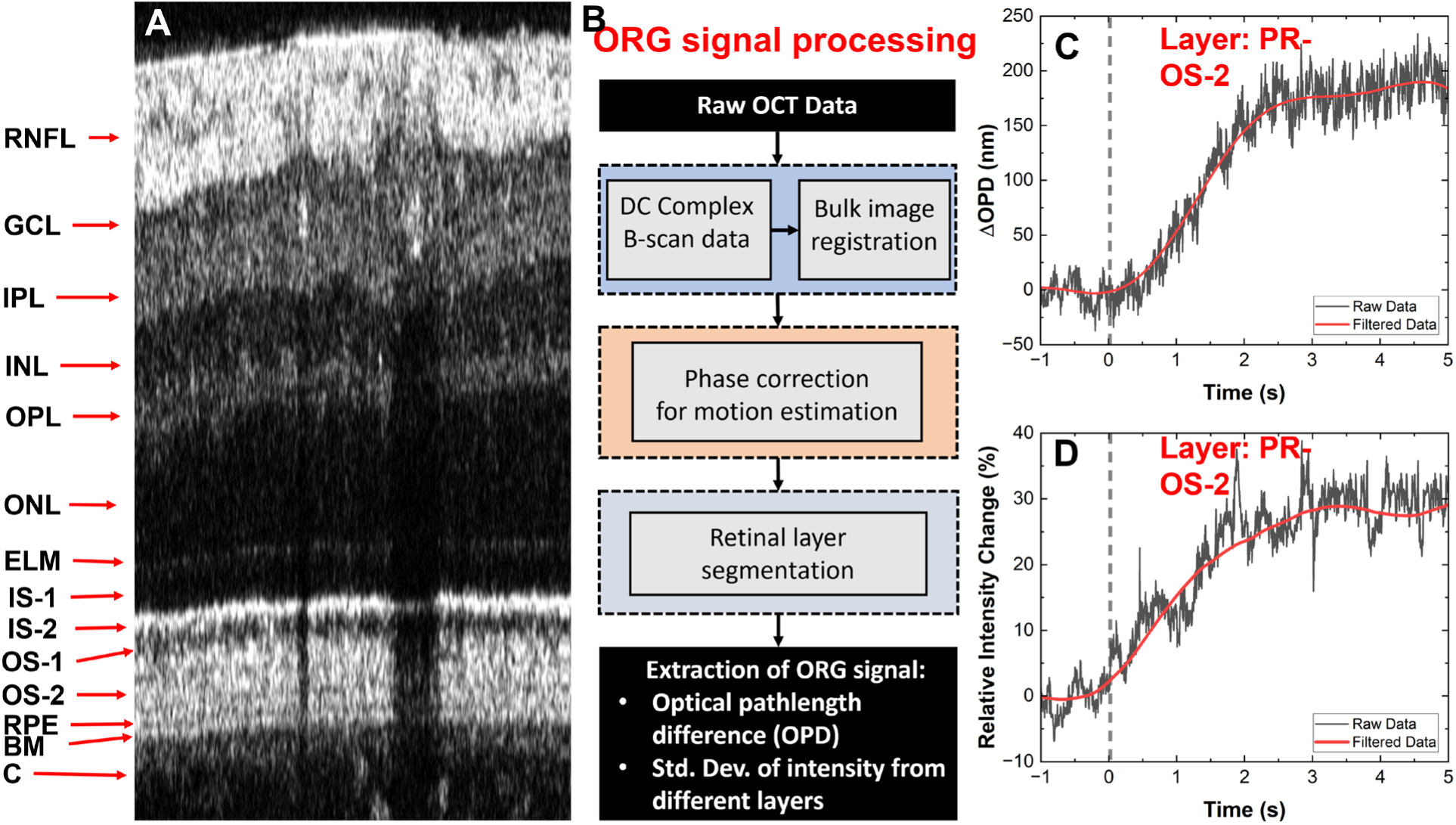
Protocol for extracting ORG traces from the repetitive B-scans. (A) A representative B-scan showing the layered structure of the human retina and cross-sections of large blood vessels in the RNFL. (B) Flow chart of the image processing steps to extract the optical path length difference (OPD) and intensity ORG traces from different retinal layers. Example ORG traces: OPD (C) and intensity (D) extracted from the OS-2 band of the photoreceptors layer.

Representative OPD and intensity traces extracted from the photoreceptors OS-2 layer are shown in Figures 3C and 3D, respectively. The high frequency data (black line) resulting from the OCT speckle pattern and periodic modulations related to breathing and heart rate were filtered out using a combination of filtering functions, in order to generate the smooth time traces (red curves) in Figures 3C and 3D.

A custom algorithm was also developed for quantifying vascular changes (retinal blood flow and blood vessel diameter) caused by visual stimulation. Figure 4A shows a representative B-scan, where an ROI including the cross-section of a retinal blood vessel is marked with the red dashed line. Figure 4B shows a flow chart of the algorithm developed for analysis of the vascular changes. Bulk registration was used to align all B-scans for eye motion.^61^ Next, a blood vessel was manually selected from the first B-scan in the time series by looking at the shadow cast by the vessel in the deeper layers of retina as blood cells are highly reflective. The width of the shadow was correlated to the inner diameter of the blood vessel as the lumen of retinal blood vessels has significantly lower reflectivity. The blood vessel was cropped from all B-scans in the time series using a semi-automated algorithm. The blood velocity was calculated based on the phase shift between adjacent A-scans.^33, 67^ Retinal blood flow (RBF) was calculated from each B-scan using the relationship:

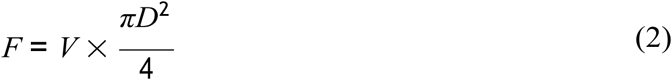

where, *F* is the retinal blood flow, *V* is the retinal blood velocity, and *D* is the diameter of the blood-vessel cross-section. Therefore, the blood velocity and blood vessel area were used to calculate RBF and track visually-evoked transient RBF changes over time.

**Fig 4.**
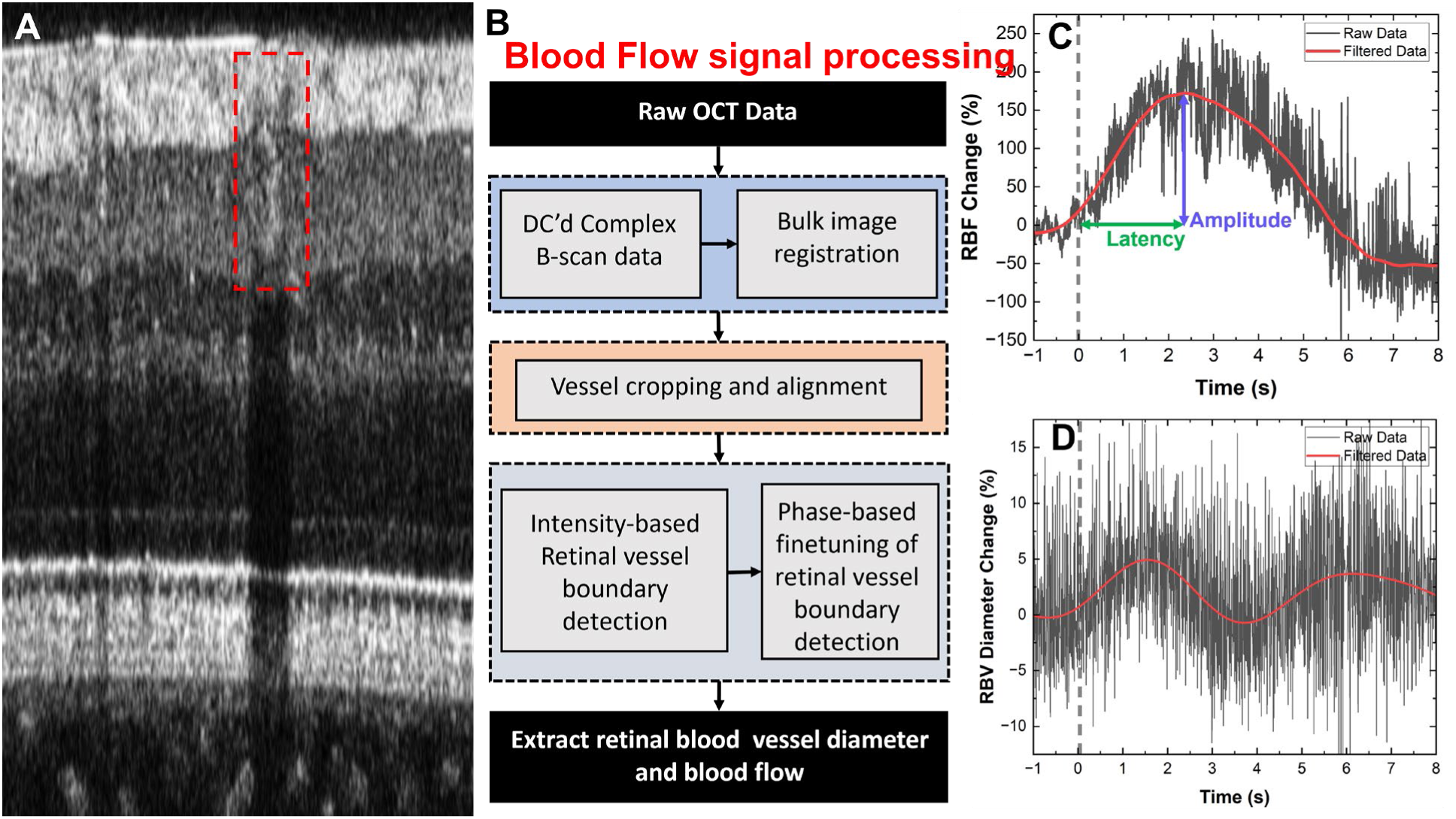
Protocol for measuring transient changes in the retinal blood vessels. (A) A representative retinal B-scan with the cross-section of a retinal blood vessel marked with the red dashed line. (B) Flow chart of the image processing steps to quantify dynamic vascular responses (blood flow change and dilation / constriction of retinal blood vessels) to visual stimulation. (C) Retinal blood flow change in response to 4 ms single flash white-light stimulus. (D) Corresponding blood vessel diameter change.

Representative RBF and blood vessel diameter (BVD) time traces are shown in Figures 4C and 4D. The raw data exhibited high fluctuations (black lines) because of speckle present in the OCT images. A smoothing filter with a moving window (Savitzky-Golay) was used to filter out the high frequency component of the data. Next, the filtered data was zero-padded (∼ 3×) to enable band-block filtering of the pulsatile fluctuations caused by the participant’s heart rate and to generate smooth traces (red lines) of the time-dependent changes of the RBF and BVD caused by the visual stimulation of the retina. Fractional changes in the RBF were computed based onthe following relation:

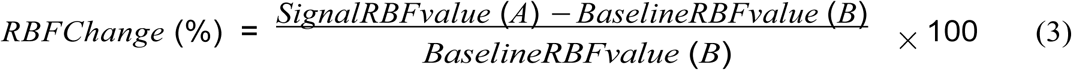

where, the signal RBF value corresponds to the RBF value obtained for a vessel in each B-scan and the baseline RBF value corresponds to the mean RBF value during the pre-stimulus period.

Similar analysis was conducted to compute fractional changes for BVD response in the axial and lateral directions:

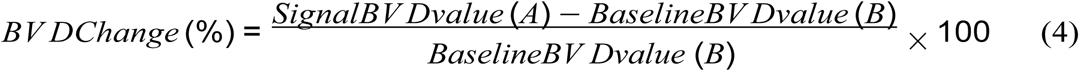

Here, the signal BVD value represents to the value of cross-sectional diameter of a blood vessel in a B-scan and the baseline BVD value is determined by taking average of the vessel diameter fluctuations over pre-stimulus recording.

To quantity the RBF and BVD responses to visual stimulation, we employed 2 metrics: ‘Amplitude’, measured relative to the baseline (pre-stimulus data) and ‘Latency’ measured from theonset of the visual stimulus to the time of the peak response (Figure 4C).

## 3 Results

Figure 5 shows representative cross-sectional and *en face* morphological images of the healthy human retina. The high axial OCT resolution allowed for clear visualization of the Bruch’s membrane (Figure 5A, green arrow), the retinal pigmented epithelium (RPE, Figure 5A, orange arrow), the posterior end of the photoreceptors outer segment (OS-2, Figure 5A, blue arrow), that contains RPE micro-villi rich in melanin granules, responsible for the high reflectivity of this sub-layer, reflections from cone tips in the anterior part of the photoreceptor’s outer segment (Figure 5A, red arrows) and the highly reflective posterior end of the photoreceptors inner segment (IS-2, Figure 5A, white arrow), that contains densely packed mitochondria. The high axial resolution also allows for visualization of three distinct sub-bands in the retinal inner plexiform layer (IPL, Figure 5B, yellow arrow) that were previously observed with research-grade visible light OCT technology with comparable axial resolution.^27^

**Fig 5.**
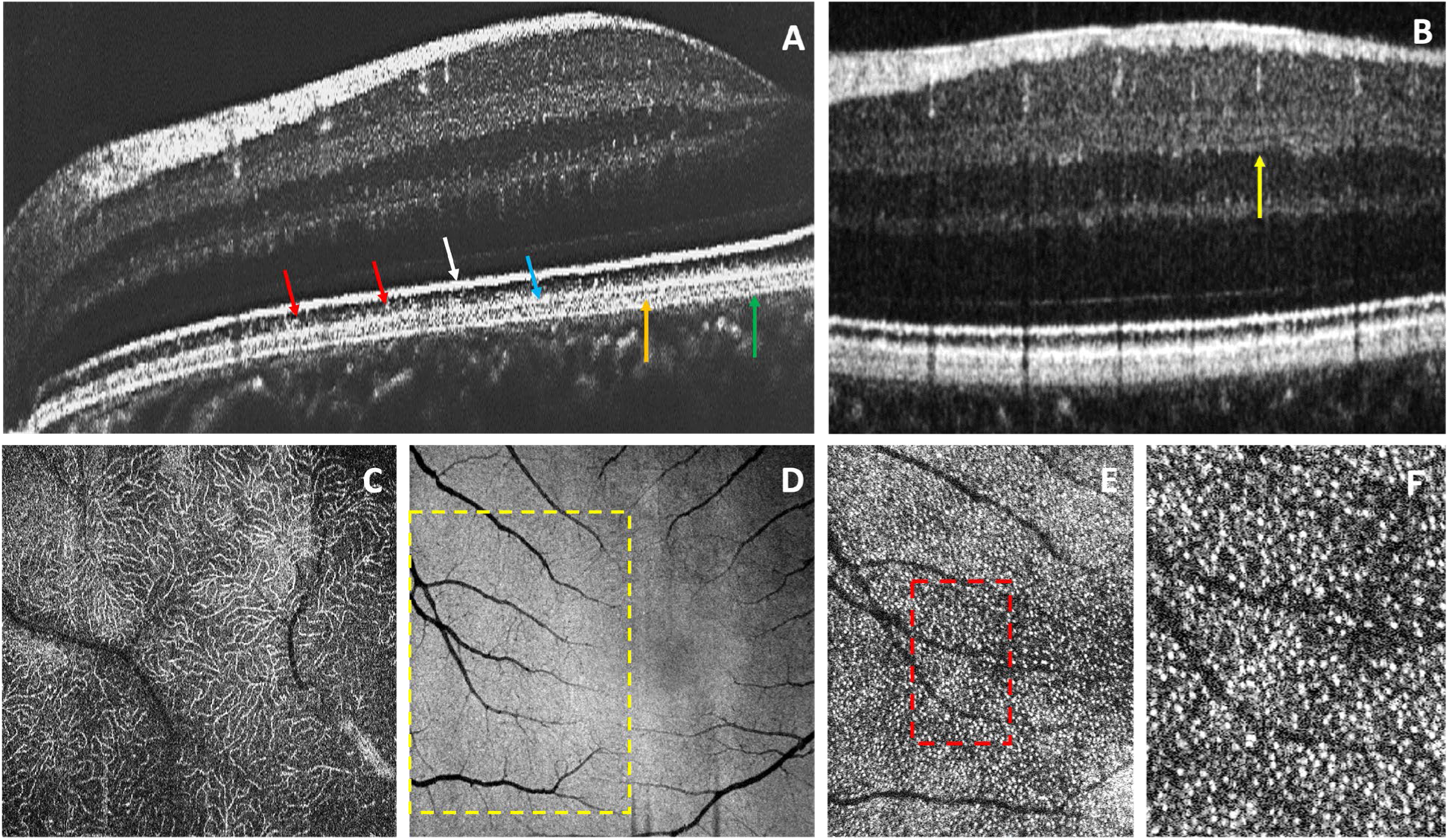
OCT images of the healthy human retina. The high spatial resolution is sufficient to resolve the Bruch’s membrane (green arrow), the RPE (orange arrow), the photoreceptors OS-2 layer (blue arrow) and reflections from individual cone OS tips (red arrow) (A), the 3 sub-layers in the retinal IPL (B) and the intricate capillary network in the retinal OPL (C). *En face* view of the photoreceptors IS/OS junction (D). Magnified views of the photoreceptors mosaic, where tips of individual cones are resolved (white dots in E and F).

Figure 5C shows an *en face* view of the microvasculature at the inner plexiform layer (IPL) of the retina. Figure 5D shows an *en face*, maximum intensity projection image of the photoreceptors IS/OS junction. Figures 5E and 5F show magnified views of a ROI located a few degrees away from the center of the fovea, where reflections from cones can be observed (highly reflective white dots).

Figure 6B shows intensity profiles of the human retina generated from a time series of B-scans (single representative B-scan from the series is shown in Figure 6A) for selected time points before, during and after the single flash stimulus. The intensity profiles were aligned with respect to each other, using the peak corresponding to the photoreceptors IS-2 layer as a reference point. A simplified pictorial representation of the retina’s cellular pathway for generation andtransmission of visional information is also shown in Figure 6A. The figure shows clearly that visual stimulation causes time dependent increase and decrease in the intensity of different retinal layers, as well as temporary swelling in some of the retinal layers, that results in shifts of the intensity peaks corresponding to those retinal layers (OPD changes).

**Fig 6.**
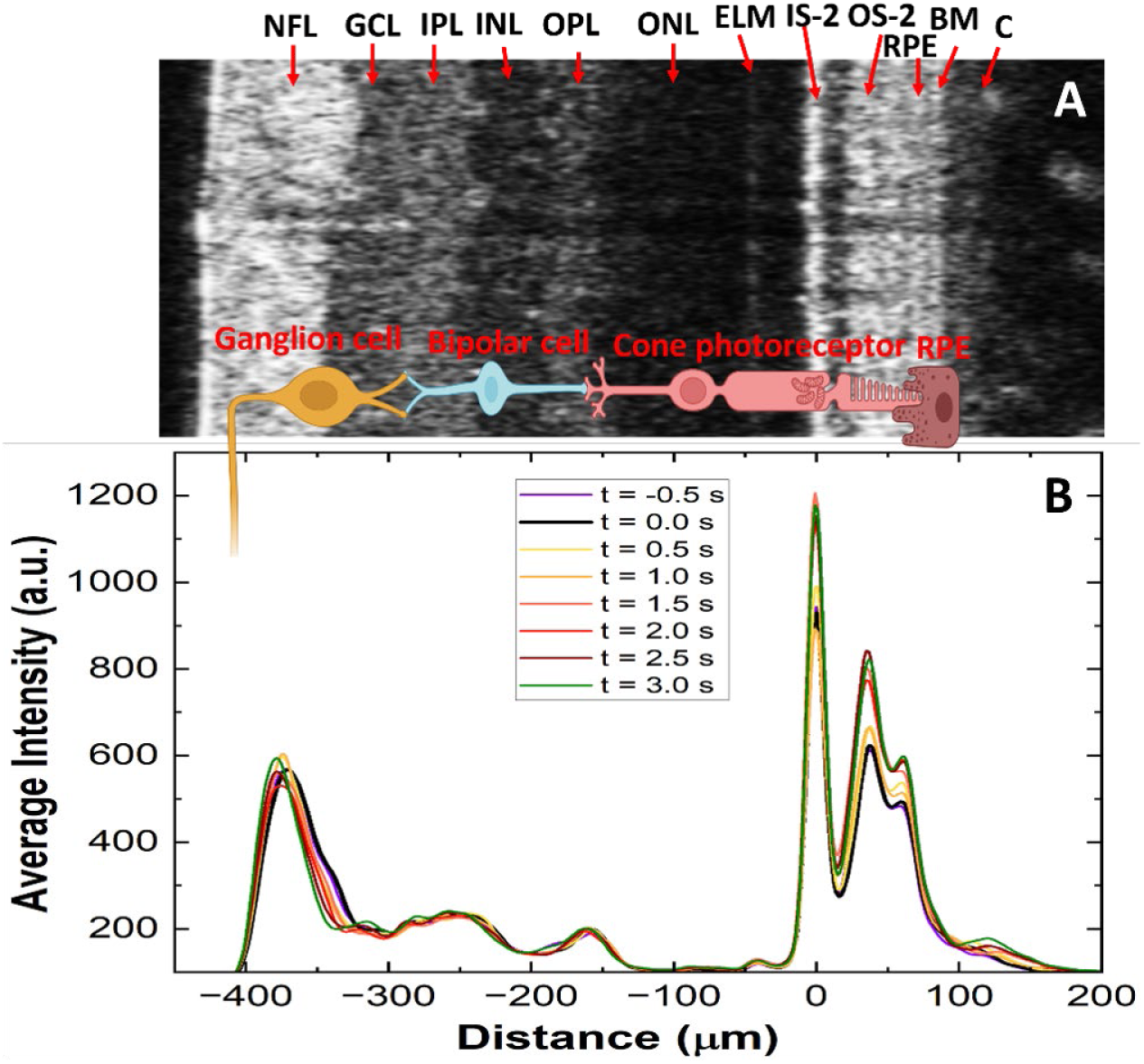
(A) Representative B-scan of the healthy human retina from a functional OCT time series with layers labeled. (B) Retina intensity profiles extracted from the series of B-scans for selected time points before, during and after application of the visual stimulus. Pictorial representation of the cellular pathway for generation of visual information.

Figure 7 shows a summary of the retinal response to visual stimulation (4 ms duration and 43.4*cd.s/m*^2^ intensity) in terms of physiological changes in the retinal neurons measured with OCT (intensity and OPD changes) and ERG, and vascular changes (transient changes in the RBF and BVD). Figure 7A shows phase-based optical path length changes in the photoreceptors OS-2 band (red line). The photoreceptors show an expansion in two steps, reaching a value of ∼ 220 nm at ∼ 5 seconds after the stimulus onset. The black line shows the OPD changes during baseline (dark recording).

**Fig 7.**
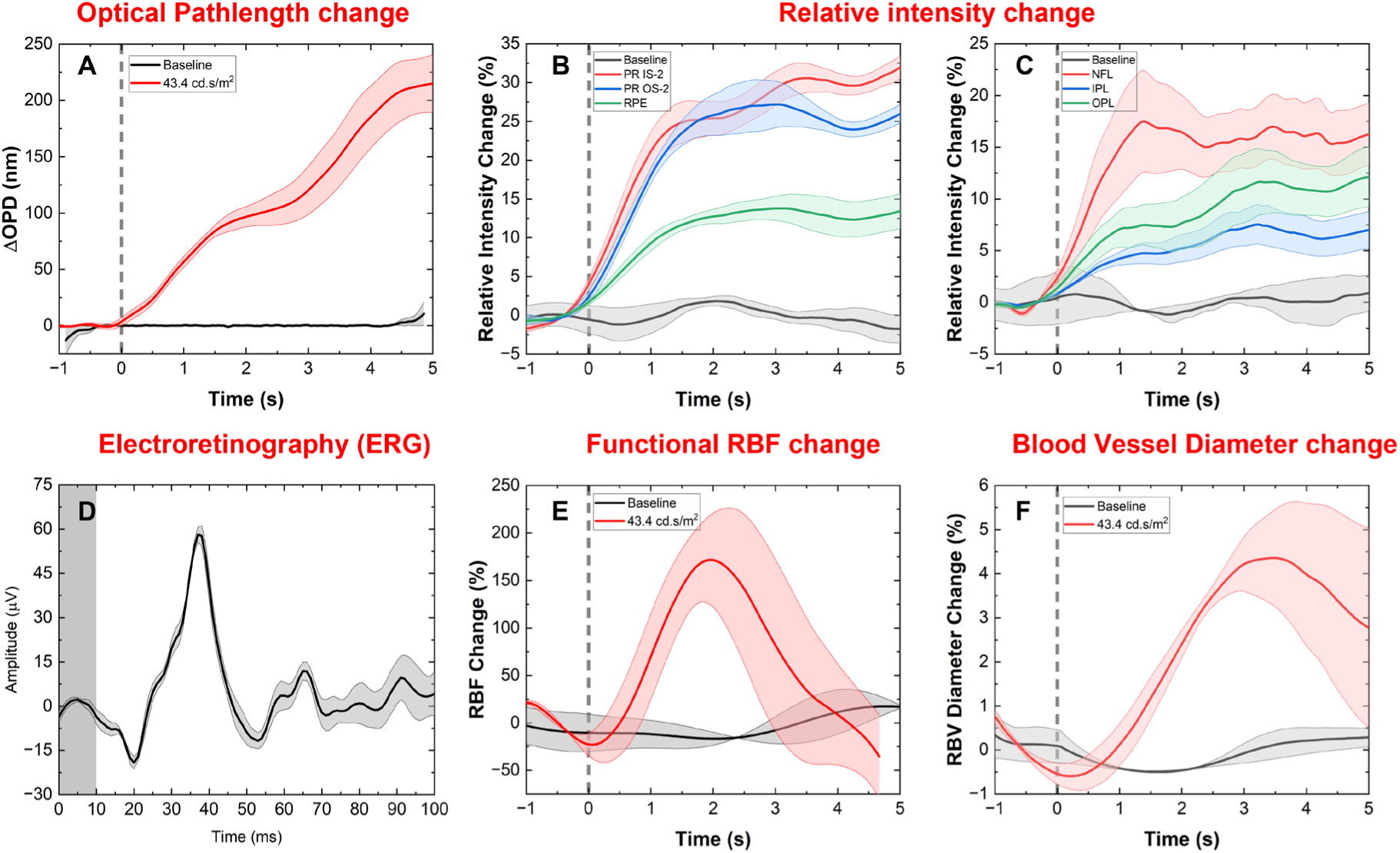
Summary of the healthy human retina response to visual stimulation (white light, single flash, 4 ms duration and 43.4cd.s/m^2^ intensity) in terms of physiological changes in the retinal neurons measured with OCT (intensity and OPD changes) and ERG, and vascular changes (transient changes in the RBF and BVD). The block line represents an average of at least 3 measurements from one subject, and the shaded area represents the standard error of the measurements. (A) Optical pathlength changes from PR OS-2 layer. Intensity changes measured from retinal layers located in the posterior (B) and anterior (C) retina. (D) Representative ERG recording. Visually evoked changes in the retinal blood flow (E) and blood vessel diameter (F).

Figures 7B and 7C show standardized intensity-based changes in the retinal layers. Figure 7B shows the intensity changes in the outer retina, specifically the IS-2 (red curve), the OS-2 (blue curve), and the RPE (green curve). The intensity changes observed in the IS-2 and the OS-2 layers are similar in magnitude (∼ 30%), however the peak intensity change for the OS-2 layer occurs a few milliseconds earlier than the change in the IS-2 layer. The intensity change in the RPE layer is of significantly lower magnitude and reaches a plateau almost simultaneously with the change observed in the OS-2 layer. The black curve shows the baseline (dark recording) response of the PR OS-2 layer.

Figure 7C shows intensity changes in the NFL (red curve), the IPL (blue curve) and the OPL (green curve). The NFL shows the highest change in intensity amongst these retinal layers, with a maximum change of *sim*18% plateauing at ∼ 1.5 seconds after the stimulus onset. The IPL and the OPL show an intensity change of ∼ 5% and ∼ 10% respectively, ∼ 1 second after the stimulus onset. The black curve shows the baseline (dark recording) response of the IPL layer.

Figure 7D shows a representative single flash ERG trace acquired for a stimulus intensity of 3.44*cd.s/m*^2^. The negative a-wave peaks at ∼ 18 ms after the flash onset, while the positive b-wave peaks at ∼ 35 ms after the flash onset. The gray rectangle in the plot shows the timing and the duration of the white light, single flash stimulus.

Figure 7E shows representative RBF traces from dark recordings (no flash, black curve) and a single flash recording with intensity of 43.4*cd.s/m*^2^ (red curve). The black curve in the figure shows the blood flow response in the absence of visual stimulation. On an average an RBF change of *<* 20% is present in the absence of visual stimulation. However, a change of as high as 200% is observed with a single flash of 43.4*cd.s/m*^2^ intensity (red curve), which peaked at ∼ 2 seconds after the flash onset. The RBF changes are transient and the flow returns to baseline within ∼ 5 seconds from the stimulus onset.

Figure 7F shows representative changes in retinal blood vessel diameter in response to a single-flash stimulus of 43.4*cd.s/m*^2^ intensity. The black curve shows the response in the absence of stimulation, where a slight pulsatile behavior is be observed. However, with the onset of stimula-tion, the vessel dilates to ∼ 4.5% reaching a maximum value at ∼ 3 seconds after the onset of the flash (red curve).

Next, we explored the dependence of the retinal response to visual stimulation of different intensity by varying the intensity of the 4 ms single white-light flash stepwise in the range 0 − 43.4*cd.s/m*^2^. Figure 8 shows representative results for 3 stimulus intensity values: 1.4*cd.s/m*^2^, 14.5*cd.s/m*^2^ and 43.4*cd.s/m*^2^. The ERG recordings (Figure 8A) showed progressive increase of the amplitude of the a-wave and b-wave, as well as small changes in the peak latency of the b-wave. The retinal blood flow (Figure 8B) showed significant increase in the peak magnitude of the transient blood flow changes with progressive increase of the stimulus intensity. The peak latency decreased initially between recordings with 1.4*cd.s/m*^2^ and 14.5*cd.s/m*^2^ stimulus intensities, and plateaued for flash intensity of 43.4*cd.s/m*^2^. Similar non-linear behavior of the RBF latency was observed by our research group in retinal veins for measurements conducted at the periphery of the optic nerve head in a different study.^68^ The recovery time of the RBF baseline is also affected by the intensity of the visual stimulus and is longer for the highest intensity of the flash. The retinal blood vessel diameter (RBVD) (Figure 8C) showed similar behavior to the RBF. The peak magnitude of the vasodilation increased progressively while the peak latency decreased with the increase of the stimulus intensity.

**Fig 8.**
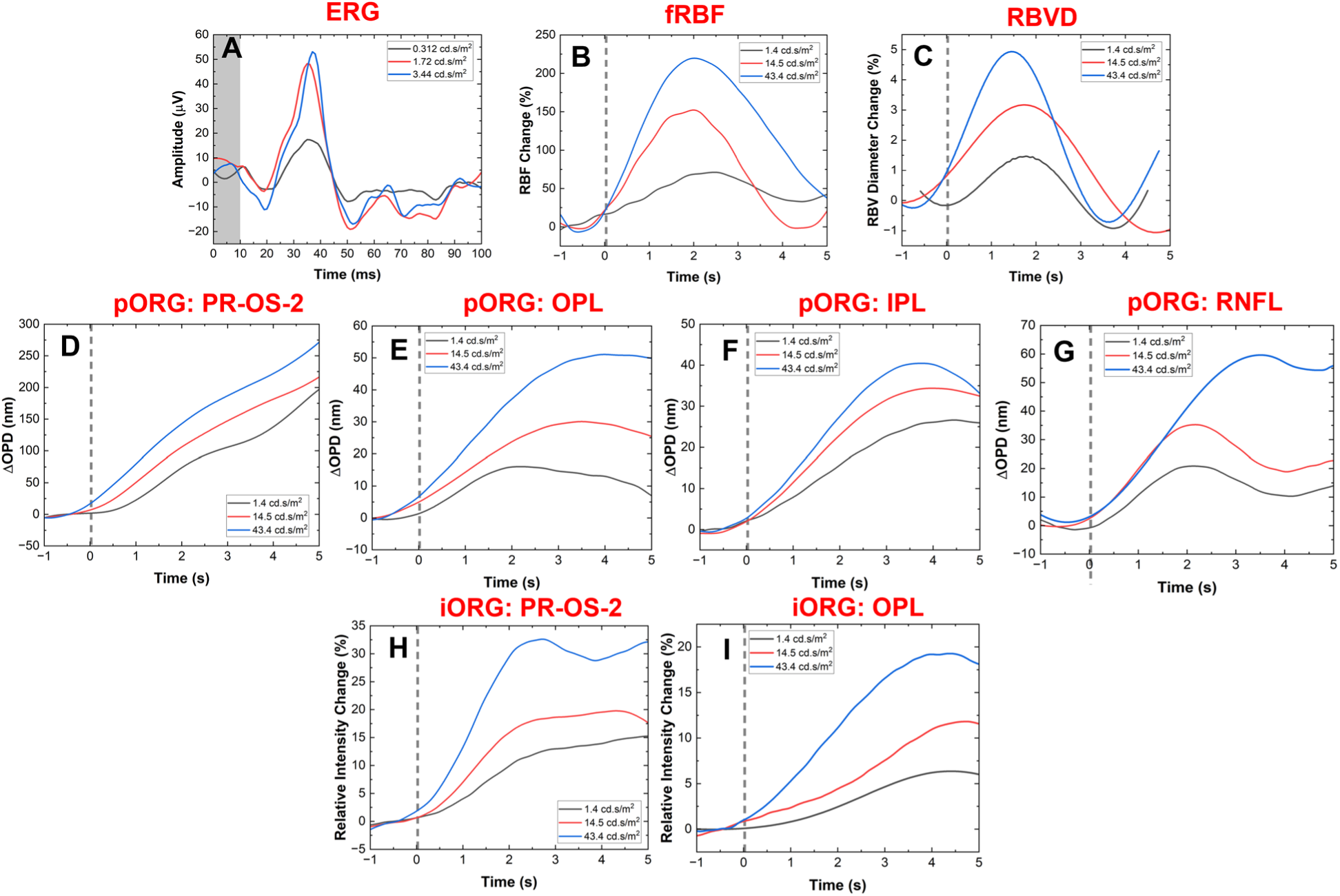
Different responses at different intensities of single-flash stimuli. (A) ERG traces; (B) RBF changes; (C) BVD changes; (D) and (H) Optical path length and intensity changes in the photoreceptor OS-2 layer; (E) and (I) Optical path length and intensity changes in the outer plexiform layer; (F) Optical path length changes in the inner plexiform layer; and (G) Optical path length changes in the retinal nerve fiber layer.

Figures 8D and 8H show the phase-based (optical pathlength difference – OPD) and the intensity-based ORG responses for the photoreceptors OS-2 layer respectively. The OPD traces showed almost linear increase over the 5 seconds time with the onset of the flash, and the response was proportional with the increase of the flash intensity, reaching a peak magnitude of ∼ 300 nm for xthe strongest flash. While the magnitude of the intensity-based ORG responses also scaled with the increase of the flash intensity, it plateaued at ∼ 2.5 seconds relative to the flash onset, reaching a peak magnitude of ∼ 30% relative to the baseline (dark recording).

Figures 8E and 8I show the phase-based (OPD) responses and intensity-based ORGresponses measured from the retinal outer plexiform layer (OPL) respectively. The optical path-length increase in this layer is almost 10 times smaller in magnitude compared to the response measured from the photoreceptors OS-2 layer, reaching peak magnitudes of ∼ 20 nm, 30 nm and 50 nm for the 3 flash intensity values respectively. The peak latency relative to the onset of the flash also increased with the intensity of the flash in the range of 2s – 4s. For the lower intensity flashes, the response showed tendency for return to baseline, while the response to the strongest flash (43.4*cd.s/m*^2^) plateaued after 4 seconds relative to the flash onset. The intensity-based responses measured for the lower intensity flashes showed transient responses of peak magnitudes of ∼ 5% and ∼ 12% and peak latencies of ∼ 1 second and ∼ 2 seconds respectively. The response to the strongest flash plateaued at ∼ 20% and ∼ 3.5 seconds latency relative to the flash onset.

Figure 8F shows the phase-based (OPD) responses measured from the retinal inner plexiform layer (IPL). The OPD traces showed progressive increase of the peak magnitude from ∼ 20 nm to x∼ 40 nm with increase of the flash intensity, while the peak latency decreased from ∼ 4.3 seconds to ∼ 3.5 seconds respectively. All of the traces showed tendency to return to baseline.

Figure 8G shows the phase-based (OPD) responses and intensity-based ORG responses measured from the retinal nerve fiber layer (NFL). For the lowest stimulus intensity of 1.4*cd.s/m*^2^, the phase-based ORG response plateaued at ∼ 20 nm, with ∼ 2 second peak latency. The OPD response to the stimulus intensity of 14.5*cd.s/m*^2^, peaked at ∼ 30 nm with ∼ 2 second peak latency. The strongest flash resulted in almost linear OPD change that peaked at ∼ 60 nm with ∼ 3.5 second peak latency.

## 4 Discussion

The novel OCT+ERG modality (imaging technology and image processing algorithms) presented here allows for simultaneous imaging of the human retina morphology and probing of retinal neuronal and blood flow responses to visual stimulation with high spatial and temporal resolution.

The OCT system is designed to operate in the same wavelength region (∼800 nm) as commercial OCT systems and has a compact fiberopic design. The high axial resolution (1.7*µm*) is sufficient to resolve the multi-layered structure of the posterior human retina (Figure 3A), including the RPE and the Bruch’s membrane, as well as the triple sub-bands of the inner plexiform layer (Figure 5B, yellow arrow) that have been observed in the past with research-grade visible light (VIS-OCT) technology of similar axial resolution.^27^ The system’s lateral resolution is sufficient to resolve reflections from the tips of individual cone photoreceptors located a few degrees away from the fovea (white dots in Figures 5E and 5F), as well as the fine capillary network of the outer plexiform layer (Figure 5C).

Integration of the OCT system with a commercial ERG technology serves severalpurposes:

1. Synchronous ORG and ERG recording that allow clinicians access to retinal morphological, functional and vascular data that is acquired simultaneously;
2. Easy, flexible control of the visual stimulus and utilization of clinically approved ERG protocols for retinal stimulation

As the ORG research field is still relatively new, simultaneously recorded ERG traces can serve as a “gold standard” to validate the ORG recordings. Note that while the current design of the visual stimulator allows for flexibility of choice in terms of timing, duration, intensity, frequency pattern and color of the visual stimulus, it can only generate a wide-field, Maxwellian view illumination of the retina. It also does not provide a fixation pattern for the imaged eye. Future designs of the visual stimulator will include fixation targets for the imaged eye, as well as ability to project spatial patterns of different sizes on the surface of the retina.

Results from this pilot study demonstrate that the OCT technology and image processing algorithms allow for measurement of both phase-based OPD changes and intensity-based changes from all major retinal layers, as well as transient changes in the retinal blood flow caused by visual stimulation of the retina. The simultaneous measurement of both neuronal responses andtransient changes in the blood flow and vasodilation of local blood vessels allows for conducing of non-invasive studies of neurovascular coupling in the human retina. Data from this pilot study shows that in general both the phase-based OPD and intensity-based responses from retinal layers populated by different types of retinal neurons, to visual stimulation increase in magnitude relative to the baseline (dark recordings) with increase of the stimulus intensity, while the latency of the peak responses decreases or increases with increase of the stimulus intensity depending on the retinal layer type.

The largest phase-based OPD changes were observed in the OS-2 sublayer of photoreceptors, which is consistent with the results of rod response-dominated animal and human studies and can be explained by the transient water influx following the phototransduction process of rhodopsin located in the outer segments of photoreceptors.^69, 70^ The OS-2 layer also shows the largest intensity change (∼ 30%) which can be associated with different factors:

1. temporary realignment of electrical dipoles within the double lipid membranes that form the OS layer, during the phototransduction process, which results in transient changes of the local refractive index;
2. conformational changes in the opsin protein can alter the scattering profile of the rhodopsin molecules in the OS layer;
3. water influx that can cause transient changes in the reflectivity profile of the OS layer.

Phase-based OPD and intensity-based changes of almost 10× smaller magnitude were observed from the outer and inner plexiform layers of the human retina in response to the visual stimulation (Figures 7 and 8). Since these retinal layers contain dense capillary networks, these changes are most likely related to transient increase in the retinal blood flow (retinal blood cells scatter light strongly) and vasodilation of the retinal capillaries. Furthermore, the inner plexiform layer consists of axons of bipolar, horizontal and amacrine cells and dendrites of the retinal ganglion cells. The increase in synaptic activity caused by the visual stimulus can cause transient, local osmotic changes in the IPL that can be registered with OCT as OPD changes. These results agree well with a previous study on OPD changes in the IPL measured by Pfaffle *et.* ^52^ with a FF-OCT system

The retinal NFL is composed of unmyelinated axons of the ganglion cells that conduct action potentials in order to transmit information from the retina to the brain. The transient increase in the NFL reflectivity measured with OCT during and immediately following the visual stimulation (Figure 8K) can most likely be explained by transient increase in the magnitude and frequency of the action potentials, which will alter the local electrical field and therefore the local refractive index of the ganglion cell axons. These changes will result in measurable changes of light reflected and scattered by the NFL.^71^ The phase-based OPD changes measured from the NFL (Figure 8G) in response to the visual stimulation are most likely due to transient swelling of the Müller glial cells, the synapses of which are located in-between NFL fibers.

The visual stimulation of the human retina caused transient changes both in the local blood flow and the vasodilation of the retinal blood vessels (Figures 8B and 8C respectively). These changes can be explained with the increased metabolic activity of retinal neurons during and immediately after visual stimulation which increases the demand for oxygen and nutrition delivered by the retinal blood network.

While we demonstrated that the temporal resolution of the OCT system is sufficient to measure transient changes in the neuronal activity (ORG recordings) and blood vasculature (RBF and RBVD) caused by visual stimulation, 10 second long recordings pose difficulties for both the imaged subjects (keeping the eye fixated and open) and the image processing algorithms (correction of eye motion artifacts). In this study, subjects tended to blink and lose fixation after ∼ 7 sec-onds, which limited the post-stimulus period to only 5 seconds and prevented exploration of the recovery process of the ORG and RBF traces back to baseline. Eye motion artifacts arising from involuntary eye motion (heart rate, breathing rate, ocular muscles twitching) also posed challenges to the filtering process of the ORG and RBF traces and caused the initial increase of the ORG and RBF changes to appear occurring slightly earlier than the stimulus onset. The limited speed of the camera in the current design of the OCT system also prevented measurement of the initial fast negative OPD response from the OS-2 layer, corresponding to the ERG a-wave, that was observed by other research groups with OCT technology with parallel detection such as SS-OCT,^31, 51, 53^ Full-Field OCT,^50^ multiple detection channels^30^ and Line-Field OCT.^54, 55^

## 5 Conclusions

In summary, we have developed a combined OCT+ERG system for in-vivo, simultaneous high -resolution imaging of the human retina structure and measurement of fast, transient changes in the retinal neuronal function (ORG and ERG) and the local retinal blood flow (Doppler OCT) induced by visual stimulation. Both optical pathlength- and intensity-based ORG signals were acquired from the major retinal layers (nerve fiber layer, plexiform layers, photoreceptor inner and outer segments and the retinal pigmented epithelium). Data acquired in this study provides insight to the neurovascular coupling in the human retina. The novel OCT+ERG technology and OCT image processing algorithms presented here can serve as valuable research tools in future biomedical and clinical studies of potentially blinding neurodegenerative retinal diseases and brain neurodegenerative conditions that are expressed in the retina.

## Code, Data, and Materials Availability

Data underlying the results presented in this paper are not publicly available at this time however may be obtained from the corresponding author upon reasonable request.

## Acknowledgments

Funding for this project was provided by the National Science and Engineering Counsel of Canada (NSERC RGPIN-2020-06308), the Canadian Institutes of Health Research (202309PJT-505682-MPI-CENA-149275) and the University of Waterloo Center for Bioengineering and Biotechnology (CBB Seed Fund, Roud 6) and the University of Waterloo Research Incentive Fund. We acknowledge James Simmons’ assistance with the development of the semi-automatic code for alignment and cropping of the retinal blood vessels.

